# Computer-assisted *segmentation-free* analysis for long-term assessment of flagellar beating in free swimming human spermatozoa: the role of emptying calcium stores

**DOI:** 10.1101/2020.07.15.204461

**Authors:** G. Corkidi, P. Hernández-Herrera, F. Montoya, H. Gadêlha, A. Darszon

**Affiliations:** Laboratorio de Imágenes y Visión por Computadora, Departamento de Ingenería Celular y Biocatálisis, Instituto de Biotecnología, UNAM, Cuernavaca, Mexico; Department of Engineering Mathematics & Bristol Robotics Laboratory, University of Bristol, UK; Departamento de Genética del Desarrollo y Fisiología Molecular, Instituto de Biotecnología, UNAM, Cuernavaca, Mexico

## Abstract

Microorganisms perform complex tasks and achieve biological function over long periods of time. Human spermatozoa are the archetype of long-time self-organizing transport in nature, and critical for reproductive success. They utilize the coordinated beating of their flagellum to swim long distances within the female reproductive tract in order to find and fertilize the egg. However, to this date, the long-time coordination of the sperm flagellum beating, or indeed other flagellated microorganisms, remain elusive due to limitations in microscope imaging and flagellar tracking techniques. Here, we present a novel methodology based on local orientation and isotropy of bio-image to obtain long-term kinematic and physiological parameters of individual free-swimming spermatozoa without requiring image segmentation (thresholding). This segmentation-free method allows for immediate evaluation of frequency and amplitude of the flagellar beat and head kinematic parameters for a duration of 9.2 min (limited by the camera’s internal memory), corresponding to hundreds of thousands of flagellar beat cycles. We demonstrate the powerful use of this technique by examining how the release of Ca^2+^ from internal stores alters flagellar beating in the long-term, as Ca^2+^ is an essential regulator of flagellar beating. We report that increasing intracellular Ca^2+^ affected head and flagellar beat frequencies and amplitudes nonlinearly over long periods of time, commensurate with physiological conditions. The simplicity and robustness of the method may offer an advantageous alternative to segmentation-based methods, and allows for straightforward generalization to other bio-imaging applications, such as *chlamydomonas*, trypanosomes, *c. elegans,* or indeed flagellated microorganisms and slender-body organisms. Thus, this method may appeal to distant communities away from sperm biology.

## Introduction

The main objective of a spermatozoon is to fertilize the female gamete. To achieve this, the mammalian sperm, which measures ~50 microns, navigates through an approximately 10-centimeter-long female reproductive tract to find the egg. With an average swimming speed ranging between 35-50 μm/s in viscous physiological media (Milligan et al., 1980; Smith et al., 2009), human spermatozoa would take approximately 100 min to cover this distance. During this journey, a spermatozoon will undergo several physiological changes due to alterations in ionic composition, flows within the female genital tract, and signals from the surrounding cells, some of which have been argued to induce chemotaxis (reviewed in Darszon et al., 2011, 2020). It is thus natural to expect that during this relatively long time, spermatozoa need to adapt their behavior to overcome the ever changing conditions and obstacles of the reproductive tract (Gaffney et al., 2011). For example, conditions in aqueous viscosity that *in vitro* promote the maturational process that occurs in the female tract allowing mammalian sperm to become able to fertilize the egg, known as capacitation, induce flagellar beat changes from high frequency, low amplitude, symmetrical flagellar beating to a vigorous, highly asymmetric one displaying deep flagellar bends, increased midpiece/proximal flagellar curving, and pronounced head lateral movements (reviewed in Stival et al., 2016). This latter mode of flagellar beating is named hyperactivation and is necessary for fertilization (Chang & Suarez, 2011). Its characteristics can vary among species and even within the same cell population (Drobnis et al., 1988; Ravaux et al., 2016), indicating possible functional switching between different sperm behaviors (Achikanu et al., 2019).

Considering the time it takes a human spermatozoon to reach the site of fertilization, it seems worthwhile to develop efficient analyzing tools that allow for very long recordings of freely swimming spermatozoa. However, the flagellar beating is quite fast, ranging from 10-25HZ, thus requiring high-speed cameras to resolve the rapid movement of the tail (>100 fps). Current Computer Assisted Semen Analysis (CASA) systems are inadequate to this task (Mortimer et al., 2015). Furthermore, the large data recordings resulting from such long periods of fast spermatozoa swimming are awkward to handle. To surmount these limitations the majority of the studies have focused on primitive head trajectories and the spermatozoa head movements for short periods of time, which are of much lower frequencies (Gallagher et al., 2018). Even so, if a better temporal resolution is available, it is still a difficult task to measure flagellar parameters.

Commonly, clinical automated analyses of spermatozoa motility are performed using CASA (Computer Assisted Sperm Analysis) (Davis & Catz, 1996) with samples constrained to two dimensional (2D) surfaces (20 microns height) and characterized only during very short periods of time e.g. 1 s and small field of views under the microscope.

Furthermore, dynamic parameters are only evaluated from the spermatozoa head detection (no direct flagellar kinematic parameters are measured). Other recently reported systems as SpermQ (Hansen et al., 2019), OpenCASA (Alquézar-Baeta et al., 2019) and Fast (Gallagher et al., 2019) require a segmentation procedure to measure either head or flagellar kinematic parameters and their overall accuracy depends on the segmentation results and imaging quality. Recently, a mathematical image analysis development allowed to accurately characterize detailed kinematics of spermatozoa flagella in 3D but only for few beat cycles (Gadêlha et al., 2020). Achikanu et al. (2019) showed that non-capacitated human spermatozoa are able to modify their swimming mode 6 times/min on average and at least once every 3.5 mins. This switching of the swimming gaits probably has evolved to contend with the changes occurring along the female genital tract.

In the present work we have endeavored to solve the above-mentioned bottlenecks by devising a *segmentation-less* image analysis that allows for the first time inspection of the dynamics of the flagellar beat over very long periods (9.2 min), corresponding to hundreds of thousands of flagellar beat cycles. The strategy is based on local orientation and isotropy features of an image (Püspöki et al., 2016) enclosing the flagellar beating of a swimming spermatozoon captured with high-speed videomicroscopy recordings. This yields numerous features from the head and the flagella movement of a single spermatozoon swimming in the microscope field for a long time. We accurately quantified the kinematics of the same spermatozoon determining four features without the need of segmentation: the (1) frequency and (2) relative amplitude of the lateral head displacement (Beat Cross Frequency BCF and Amplitude of Lateral Head displacement ALH in CASA standards), (3) the frequency and (4) relative amplitude of the flagellar beat (Flagella Frequency Beat FFB and Amplitude of Flagellar Beat AFB).

We demonstrate the powerful use of our segmentation-less technique by examining how the release of Ca^2+^ from internal stores alters flagellar beating in the long-term. Intracellular Ca^2+^ and its concentration changes have been implicated in the regulation of flagellar properties (Darszon et al., 2011; Strünker et al., 2015; Lishko and Mannowetz, 2018). In many cell systems it is known that the Ca^2+^ released from internal stores contributes significantly to the increase of intracellular Ca^2+^ concentration ([Ca^2+^]i) that regulates many cell signaling responses (Clapham, 2007; Putney, 2013). In this context, this work shows how the methodology presented here can be also used to examine how the swimming spermatozoa characteristics are modified by emptying its Ca^2+^ stores with two well-known Ca^2+^ regulators of these stores that inhibit their Ca^2+^-ATPase, thapsigargin (Andersen et al., 2015) and cyclopiazonic acid (Chang et al., 2009). The strategy permitted taking short intervals (avoiding cell damaging) of a [Ca^2+^]i sensitive dye emitted fluorescence for long-term evaluation, before and after applying these drugs to the same spermatozoon. Increasing [Ca^2+^]i in this manner affected several fold the head and flagellar beating frequencies and relative amplitude of the head beat (ALH) in a convoluted manner.

The main advantages of the proposed methodology are that they avoid the substantial effort needed to segment and track flagella showing high robustness to contend with different sources of microscope imaging, noise, light heterogeneity insensitivity and debris, as well as its fast processing time of large data sets with tens of thousands of image frames. This makes the method easily applicable to an umbrella of bio-imaging applications, such as in CASA systems for sperm analysis, from slender-body organisms to flagellated microbes, including, but not limited to, spermatozoa of any specie, *Chlamydomonas (*Polin et al., 2009), trypanosomes (Gadêlha et al.,2007) and *c. elegans (*Ding et al., 2019). Finally, the estimated parameters obtained with this methodology were compared with those estimated with segmentation-based systems observing high correlations.

## Material and Methods

### Ethical approval for the spermatozoa samples

The Bioethics committee of the Institute of Biotechnology, UNAM approved the proposed protocols for the human spermatozoa sample handling. The donors were properly informed regarding the experiments that would be performed and a Consent form was signed and agreed by each donor. All World Health Organization requirements were fulfilled in this study.

### Biological preparations and dye loading

After a minimum period of 48 hrs of sexual abstinence, healthy donors masturbated and human spermatozoa samples were collected. Highly motile spermatozoa were selected by a one hour swim-up protocol (Ham’s F-10 medium at 37 °C in an atmosphere of 5% of CO_2_ and 95% air). The collected cells were centrifuged at 3000 RPM for 5 min and resuspended at a concentration of 10^6^ cells/ml in a physiological solution that contained (mM): 120 NaCl, 4 KCl, 2 CaCl_2_, 1 MgCl_2_, 25 NaHCO_3_, 5 Glucose, 30 HEPES, 10 Lactate at pH 7.4. To measure [Ca^2+^]i, cells were incubated in a medium containing the Ca^2+^- sensitive Fluo-8-AM dye at 10 μM for 60 min and then washed with the same medium without dye once by centrifugation for 5 min at 3000 RPM to remove the remaining external dye. Thapsigargine (5 μMM) and cyclopiazonic acid (5 μMM) were administered to examine how the release of Ca^2+^ from internal stores affected the long-term motility of individual spermatozoon.

### Spermatozoa samples and statistical analysis

A total of 88 spermatozoa were analyzed for periods of up to 9.2 min (4.6 min for bright field (BF) or 9.2 min for fluorescence). Twenty two cells were recorded before and after thapsigargin and 22 for cyclopiazonic acid for kinematic analysis. Eighteen cells were used for fluorescence analysis. We performed two types of controls: a) for the same individual spermatozoon, the first 45 s without the application of thapsigargin or cyclopiazonic acid and compared with the recorded behavior after being exposed to the Ca^2+^ modulators. b) Twenty six spermatozoa were recorded without applying Ca^2+^ modulators (long term time-control) for 4.6 min (bright field). To determine the effect size and its statistical significance between spermatozoa characteristics before and after applying the Ca^2+^ modulators, we measured the Cohen’s *d* Effect Size for Anova and Kruskall Wallys tests for not normal populations (Eq. 1,2)). Cohen’s *d* Effect size can be used to compare two means and is defined as:

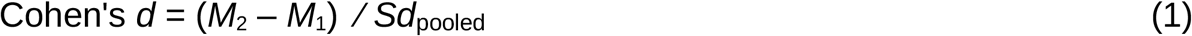

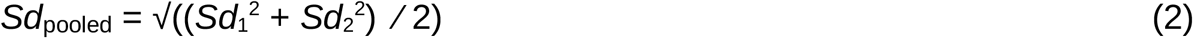

being the ratio between the difference of means before (*M*_*1*_) and after (*M*_*2*_) Ca^2+^ modulator application divided by their corresponding pooled standard deviation calculated from the standard deviations of means before and after, respectively, (*Sd_1_)*and (*Sd*_*2*_). For a d=1 or 0.5, the two group means differ by one or by a half standard deviation respectively. Cohen proposed that a small, medium or large effect size, corresponds roughly to a d= 0.2, 0.5 or 0.8 respectively (Nakagawa and Cuthill, 2007).

### Experimental setup

Experiments were performed with an Olympus IX71 (Olympus America Inc., U.S.A.) inverted optical microscope configured for bright field illumination (for head and flagellum kinematic analysis) with a LUCPlanFLN 40x/0.6 NA Objective (Olympus America Inc., U.S.A.). A UIS-2 LUMPLFLN 60X/1.00 N.A. water immersion objective (Olympus America Inc., U.S.A.) was used for fluorescence analysis. A 49011 Fluo cube Filter (Chroma Technology Corporation, USA) and a high intensity excitation LED M490D2 (Thorlabs, USA) were employed in these experiments. To keep individual spermatozoon in the field of view swimming freely in 3D we utilized a motorized x, y stage (Märzhäuser Wetzlar GmbH & Co. KG, Germany) driven by a LUDL 5000 controller (Ludl Electronic Products, Ltd., USA) in combination with the mouse pointer, using bespoke Matlab algorithms (The MathWorks, Inc., USA), from which the x,y stage position coordinates corresponding to the spermatozoa displacement were recorded. The motorized stage is not essential to image spermatozoa swimming. A manual stage can be used instead to keep individual spermatozoon in the field of view. We used a high-speed camera NAC Q1v (N ac Americas, Inc., USA) with 8 Gigabyte RAM (recording up to 4.6 min at 100 frames per s with a spatial resolution of 640 × 480 pixels for bright field experiments and 9.2 min at 50 fps for fluorescence experiments). A TMC optical table (GMP SA, Switzerland) shielded the optical system from external vibration. A temperature of 37° C was kept constant with a thermal controller TCM/CL-100 (Warner Instruments LLC, USA). Data acquisition and image analysis was conducted with an Intel® Core(TM) i7-6700 CPU @ 3.4 GHz, 32 GB RAM processor (Intel Corporation, USA).

### Image acquisition and segmentation-free analysis

#### a. Bright field and fluorescence image acquisition

Spermatozoa at a low density (~10^2^ cells/ml) were placed in a Chamlide CMB chamber having a 18 mm diameter coverslip at the bottom, which was mounted on the microscope stage and temperature controlled at 37° C. Individual spermatozoon swimming at the focal plane of the coverslip were randomly selected for analysis after preparation and dye loading (see Methods: Biological preparations and dye loading). Once a spermatozoon was selected, the microscope stage was moved to keep the spermatozoon in the field of view for the duration of the experiment. This was achieved by following the spermatozoon with a mouse pointer that controlled the microscope stage and a focus motor for 4.6 min acquiring images with the digital camera at 100 fps (for bright field) and for 9.2 min at 50 fps (for fluorescence experiments). A total of 28,000 images were acquired in this fashion independently of the illumination mode.

For fluorescence image analysis, we implemented an electronic switch based on an Arduino UNO platform (Evans, 2008) alternating between bright field illumination and fluorescence. To acquire fluorescence information during 9.2 min without bleaching or damaging spermatozoa, fluorescence was sampled for 0.2 s every 4.8 s of bright field illumination (Fig. 2). For both bright field and florescence imaging, the first 45 s were recorded as a control without the Ca^2+^ regulatory compounds. After 45 s, thapsigargin or cyclopiazonic acid were administrated with a pipette, while recording continued for a given individual spermatozoon. Control experiments were also conducted without any drug administration for 4.6 min for bright field experiments. A total of 88 free-swimming spermatozoa were recorded and tracked, 26 for time control (no drugs), 22 for thapsigargin, 22 for cyclopiazonic acid and 18 for fluorescence.

#### b. Computer-assisted segmentation-free measurements

The critical advantage of the implemented method is the *segmentation-free* principle. This means that it is not necessary to segment the flagella from the videomicroscopy files to extract kinematic information, instead, it obtained directly from raw imaging data. Additional important advantages are the insensitivity to defocusing and inhomogeneous illumination. Our method is based on the local orientation and isotropy features of an image (Schindelin et al., 2012; Püspöki et al., 2016), which can be used to extract the “dominant orientation” of a single swimming spermatozoon in the image (see details in Supporting Information). The image orientation is correlated with the angle between the sperm head and flagellum. As a result round objects and debris are automatically discarded given their mostly isotropic aspect ratios, since no dominant direction exists. Fig. 1 shows a spermatozoon in the microscope field of view and its flagella pseudo-colored according to the local image orientation (in degrees) relative to the x axis. The dominant direction of an image corresponds to the average of all directions of detected grayscale gradients in each image pixel frame.

**Fig. 1.**
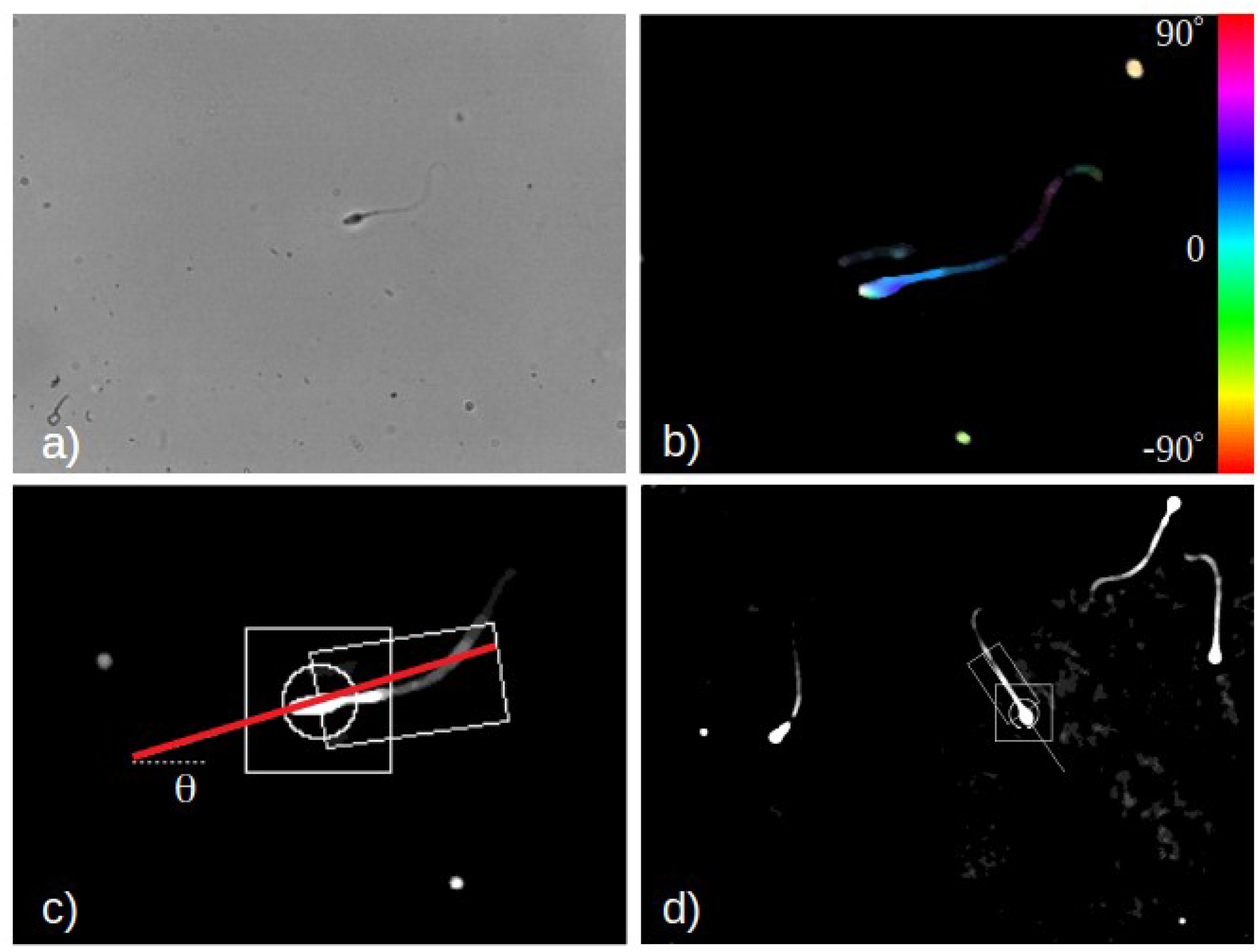
Extraction of sperm flagellum kinematics without segmentation. a) A free-swimming spermatozoon in 3D not confined in space. b) Local orientation of the image extracted from grayscale gradients (pseudocolor corresponds to the local orientation angle relative to the x-axis, see SI). c) Measuring the spermatozoon head and flagella orientation based on the local average orientation *θ (t)* of the image at time frame *t* - the square is used to follow the head in consecutive time-points, while the circle and rectangle are used to compute the sperm orientation. d) By tracking individual spermatozoon, other cells and debris are automatically ignored with this method, only the square and rectangle ROI’s are considered to obtain the motility and fluorescence measurements reported.

While the method works satisfactorily with one single cell in the field of view (for low density samples), we have modified and extended the functionality of OrientationJ (Püspöki et al., 2016) for multiple cells in the same field of view. This provides a robust dominant orientation detection avoiding the influence of artifacts and other cells when working with larger cell densities. This was achieved by tracking the spermatozoon head only and measuring the dominant orientation locally, in the vicinity of the sperm head (see details in Supporting Information). In this way, the dominant orientation of the spermatozoon image, including its flagellum, is obtained by removing any image orientation bias caused by nearby swimming cells (see Fig. 1d and Supporting Movie 1 for over than 250 s of duration). Supporting Movie 2 shows the robustness of the algorithm for a very noisy background in the image and scattered debries (see t= 180 s and 260 s). With this procedure, we extracted four different flagellar kinematic features directly from each raw video recording with durations of 4.6 and 9.2 min, respectively for bright field and fluorescence. This is described in Table 1 and also depicted in Fig. 3. Frequency and amplitude measurements were calculated directly with windowed Fast Fourier transforms of the image orientation signal measured as time advanced (see details in Supporting Information).

**Table 1.** Parameter definitions obtained with the proposed *segmentation-free* methodology (see validation of these parameters in the Section on Methodology validation).

| **<u>FREQUENCY PARAMETERS</u>** | Description |
| --- | --- |
| Beat Cross Frequency (BCF) | BCF is defined as the number of times the spermatozoon head crosses the average direction of movement (see OpenCASA, Alquézar-Baeta et al., 2019). Here we estimate BCF from the <u>first</u> frequency-peak of the Fourier transform of the orientation signal $\theta(t)$ (see Fig. 3a, b, c). |
| Flagellar Beat Frequency (FBF) | FBF is estimated from the <u>second</u> frequency peak of the orientation signal $\theta(t)$ in the Fourier transform (see Fig. 3a, b, c). |
| **<u>AMPLITUDE PARAMETERS</u>** | Description |
| Amplitude of Lateral Head Displacement (ALH) | ALH is defined by CASA as the value of the head displacement with respect to the mean trajectory (see Katz, 1991). Here we estimate ALH from the relative amplitude of the <u>first</u> frequency-peak of the Fourier transform of the orientation signal $\theta(t)$ (see Fig. 3a, b, d). |
| Amplitude of Flagellar Beat (AFB) | This parameter is defined as the ‘head centerline deviation’ D, (see Smith et al., 2009) to extract the relative amplitude of the flagellar beating. Here we estimate AFB from the relative amplitude of the <u>second</u> frequency-peak of the Fourier transform of the orientation signal $\theta(t)$ (see Fig. 3a, b, d). |

The *segmentation-free* method provides a direct measurement of the principal orientation of the image containing the spermatozoon as a whole for each time frame. As such, the orientation angle captures the combined effect of both head lateral movement and the shape of the flagellum. Thus, the method provides a time series of the instantaneous orientation angle *θ (t)* of the spermatozoon image at time *t*, relative to the x-axis along the microscope image. Head yawing and flagellar beating are extracted directly from *θ (t)* using Fourier Transform decomposition. The Fourier transform of the orientation signal *θ (t)* shows typical two frequency-peaks in the spectrum (Fig. 3b). The lower frequency-peak defines the frequency of the head yawing that occurs at a slower rate than flagellar beating frequency. The second-frequency peak in the Fourier spectrum of *θ (t)* captures the flagellar beating frequency. The associated amplitudes of the first and second-frequency peaks of the Fourier spectrum thus provides a measure of the amplitude of the oscillations of the orientation *θ (t)*, respectively, associated with the head lateral movement and flagellar waving amplitude. Note that the flagellar wave amplitude defined here is relative to the mean sperm orientation for a given instant. Four sperm kinematic parameters (table 1) are thus extracted directly from an one-dimensional signal *θ (t),* the frequency and amplitudes of both head yawing and flagellar waves, highlighting the simplicity, robustness and reduction of dimensionality of the proposed method. The sperm kinematic parameters obtained via the *segmentation-less method* have been compared with direct measurements of both head and flagellar oscillations using segmentation-based techniques reported in the literature. A strong correlation was found with the *segmentation-less* measurements reported here, as discussed in the results section.

#### c. Segmentation-free fluorescence measurements to monitor Ca^2+^ over long periods

[Ca^2+^]i was monitored via Fluo-8 fluorescence by acquiring 0.2 s samples every 4.8 s. This sampling rate allowed recording [Ca^2+^]i for long periods before and after the application of Ca^2+^ regulatory compounds without bleaching or damaging the spermatozoon. At the same time, this sampling rate is sufficiently short for long periods of Ca^2+^ evaluations, with a time resolution of 5 s, sufficient to obtain quantitative kinetic parameters from Fourier transform analysis. Fig. 2 illustrates the sampling periods and the switch between bright field and epi-fluorescence illumination.

**Fig. 2.**
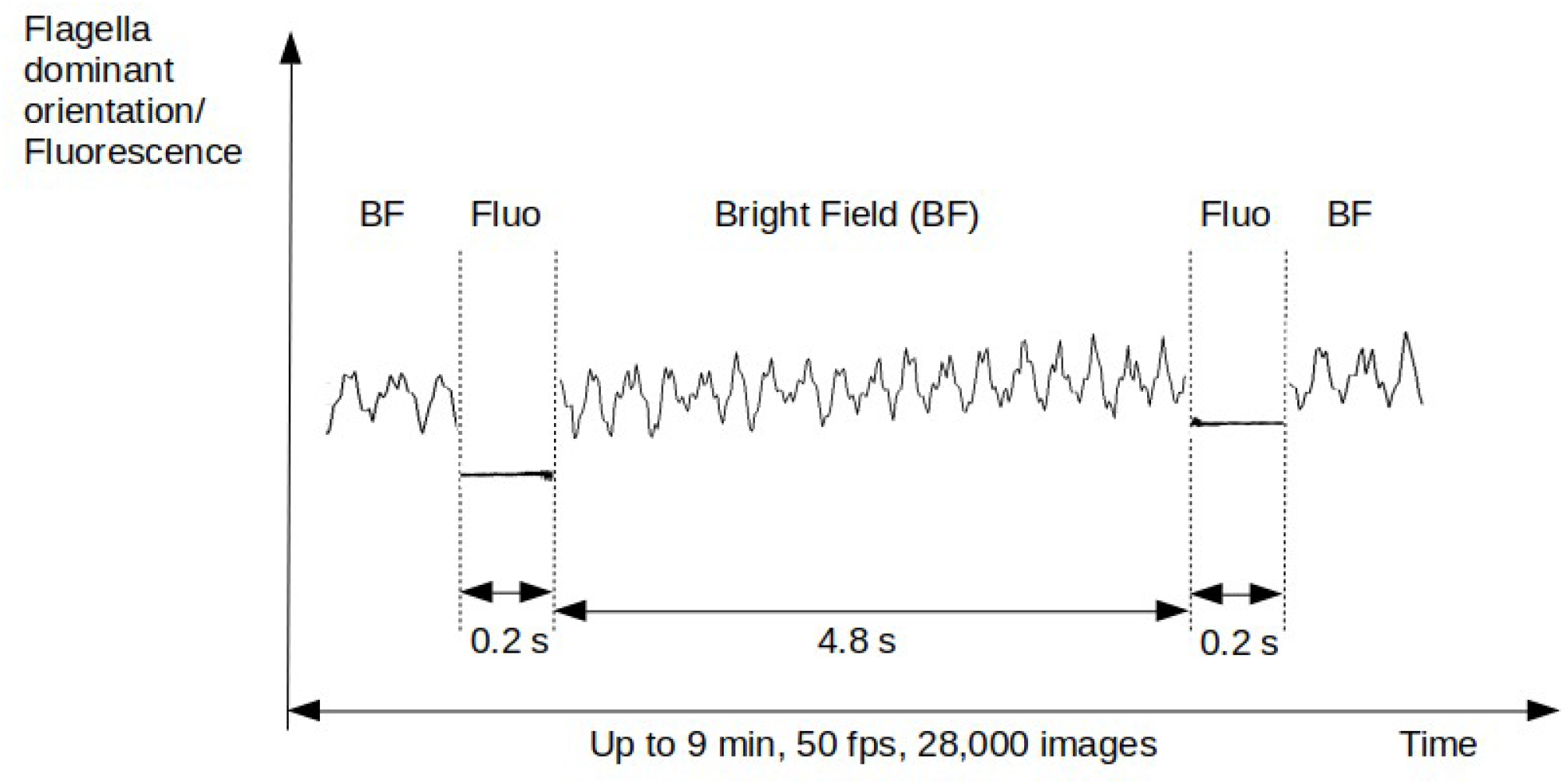
Fluorescence (Fluo) vs bright field (BF) sampling intervals. Each 4.8 s, bright field capture of the flagellar movement is switched to epi-fluorescence for 0.2 s to acquire fluorescence intensity information. For up to 9 min (at 50fps), 28,000 images are recorded with the interlaced information.

## Results

We have implemented a computer-assisted *segmentation-free* method to obtain long-term kinematic and physiological parameters for up to 9.2 min from individual free-swimming spermatozoa not requiring flagella image segmentation which is cumbersome in nature. As in many cellular systems, the release of Ca^2+^ from internal stores is well known to contribute significantly to [Ca^2+^]i homeostasis, thus driving specific signaling responses (Clapham, 2007; Putney, 2013). We demonstrate the powerful use of this technique by examining how spermatozoa swimming characteristics (head and flagellum movement) are modified by emptying the Ca^2+^ stores with two different store Ca^2+^ regulators: thapsigargin and cyclopiazonic acid. We report that increases in [Ca^2+^]i significantly affected the head and flagellar beating frequencies and relative amplitude of the head beat (ALH) in a complex dynamical manner.

Fig. 3 shows for two swimming spermatozoa (control -red- vs pharmacologically perturbed case -black-), the four extracted kinematic parameters and how [Ca^2+^]i behaves before and after applying thapsigargin. Fig. 3a shows the angle θ *(t)* rescaled by 2π, as depicted in Fig. 1c, corresponding to the dominant orientation of the sperm as it swims. Head and flagellar beat characteristics are extracted directly from θ *(t)* using Fourier transform analysis (see Methods: Computer-assisted *segmentation-free* measurements). A close-up of θ *(t)* for the small region marked with a black rectangle in Fig. 3a is shown in Fig. 3b. This unveils the characteristic two frequency-peaks of θ *(t)* associated with the head yawing and flagellar beating. Figs. 3a and c show complex behavioral changes for both head and flagella oscillations before and after the administration of thapsigargin. The sperm flagellum hyperactivates as [Ca^2+^]i is increasing and then stalls when levels reach a plateau, as indicated by yellow and blue markers in Fig. 3f and Supporting Movie 1. The positive slope of the sperm orientation angle relative to the x-axis (Fig. 3a, θ) indicates counter-clockwise rotations of the sperm flagellum as it swims (Fig. 3f). In Fig. 3a, the orientation angle is divided by 2π, thus capturing the number of turns as the orientation of the swimming direction changes during the course of the experiment. In Fig. 3a The sperm associated with the “black” curve in Fig. 3a starts roughly aligned with the x-axis at the beginning of the experiment and slowly rotates its swimming direction counter-clockwise, as also depicted in Fig 3f for its swimming trajectory. This sperm completes almost two full rotations in 4.6 min (anti-clockwise, see Fig. 3f). This highlights how swimming sperm orientation varies tremendously during long time-scales, thus correlating weakly with past swimming directions. This is indicative of intermittent dynamical beating asymmetries that are taking place over long swimming periods, thus facilitating sperm transport and increasing sperm diffusion in this time-scale.

**Fig. 3.**
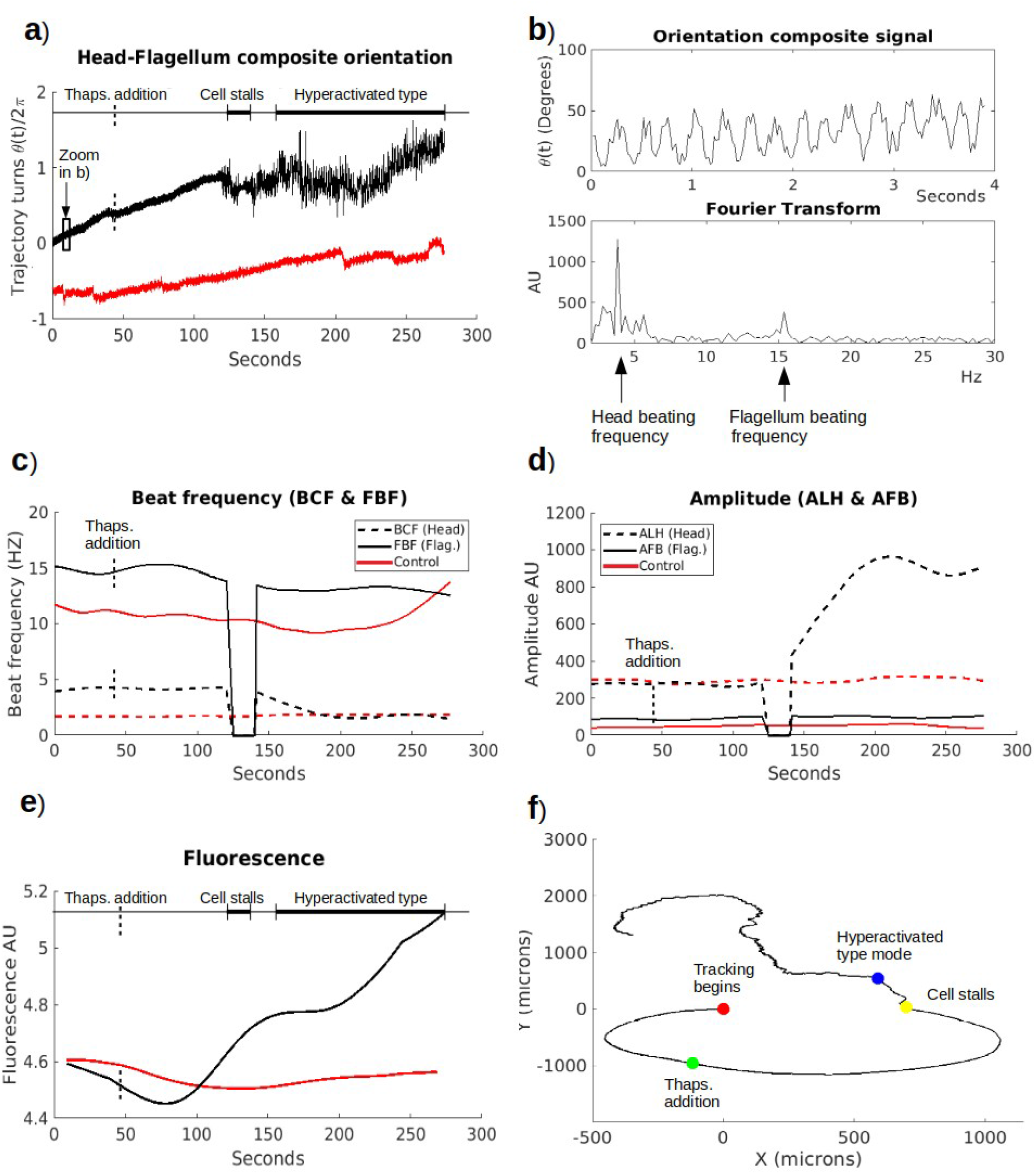
Motility parameters and [Ca^2+^]i changes for two free swimming spermatozoa (control -red- vs thapsigargin treated -black-) extracted with the *segmentation-free* method. a) Composite orientation signal θ *(t) rescaled* by 2π illustrating several stages before and after thapsigargin addition. After a significant delay from the thapsigargin addition (~75 s), the cell briefly displays a hyperactivated type mode before stopping, as the rate of [Ca^2+^]i increase slows (see 3e), before approaching the plateau. Thereafter, when [Ca^2+^]i is no longer increasing during the plateau, the sperm re-initiates motility displaying a hyperactivated type behavior which becomes more exacerbated as [Ca^2+^]i continues increasing. b) Close-up of the swimming orientation θ *(t)* in a) (top), and the Fourier transform (bottom) showing two principal peaks corresponding to head yawing and flagellar beating frequencies, defining BCF and FBF respectively (described in Table 1 and SI) depicted in c). The amplitude of the FFT spectrum providing ALH and AFB (see Table 1 and SI) are shown in d). c) Head (dashed line) and flagellar (continuous line) beating frequencies as a function of time (both decreasing up to 200 s). d) Head (dashed line) and flagellar (continuous line) beating amplitudes as a function of time (ALH increasing up to 220 s while AFB near constant). e) [Ca^2+^]i followed by Fluo-8 fluorescence intensity as a function of time (increasing for sperm thapsigargin treated). f) Trajectory of the spermatozoon thapsigargin treated with red point: tracking begins; green point: addition of thapsigargin addition; yellow point: when the flagellum stalls; blue point: the exact moment when the swimming gait changes moving away from the starting focal plane in 3D (SI movie 3).

Fig. 3c shows how the head yawing and flagellum beating frequencies decrease after injection of thapsigargin (up to 64% and 19% for the head yawing and flagellum frequency, respectively). However, this change is not immediate, it occurs after a significant time-delay (~75 s). In contrast, the control sperm depicted in red remains stable and only shows a large increase in beating frequency towards the end of the imaging period, demonstrating the potential for spontaneous self-switching, in agreement with earlier observations (Achikanu et al., 2019). Fig. 3d shows that after decreasing to zero, when the flagellum stalls, the amplitude of the thapsigargin treated sperm head lateral motion (arbitrary units) increases dramatically by 242%, and very fast, while that of the flagellum also increases, though mildly (22%), as Ca^2+^ plateaus (note that this captures the amplitude of the flagella beat relative to the head, elsewhere known as deviation, as detailed in Methods section). Fig. 3e depicts [Ca^2+^]i and show an increase of 16% after 45 s after administration of thapsigargin. This cell roughly displays a hyperactive type behavior since both head and flagellum beating frequencies decrease whereas their relative amplitudes increase, at the same time that Ca^2+^ increases and plateaus (Chang & Suarez, 2011; Ooi et al., 2014). The red curve for the control sperm in 3e shows no significant change in [Ca^2+^]i nor in swimming behavior until ~250 s when only its beating frequency increases (Fig. 3c). Fig. 3f illustrates the trajectory in the x-y plane of the pharmacologically treated sperm in Fig. 3a (black curves), where a red dot indicates the beginning of the journey and a green one the inhibitor addition. After a time delay of about a min after the thapsigargin addition, the trajectory undergoes significant changes. The sperm swims close to 2 full anti-clockwise circles during the 4.6 min, performing a total of 3850 beat cycles. This is depicted in Supporting Movie 3 which includes its evolving sperm trajectory. The sperm swimming behavior can be divided into three swimming gaits: 1) 0 – 120 s, the sperm swims in a activated progressive manner covering a long distance; 2) 120 - 140 s, after a short period of hyperactivated type mode (6 sec) the sperm abruptly stops and thereafter initiates rapid flagellar shape changes with high curvature (yellow point in Fig 3f and Supporting Movie 3), and 3) 159 – 277 s, sperm clearly switches to a hyperactivated gait (blue point in Fig. 3f and Supporting Movie 3), characterized by a decrease in flagellar beating frequency and progressivity, and an increase of its amplitude (Figs. 3c,d). At this stage the sperm swims more randomly oscillating between straight-swimming and turning periods due to vigorous whip-like flagellar movements. This is apparent in the jittery trajectory after the blue mark in Fig 3f and at 160 s in the Supporting Movie 3. It is worth noting that this sperm was actually swimming in three-dimensions, moving away from the starting focal plane. For x-y recording purposes this was compensated by refocusing the microscope as required during the experiment. Fig 3f thus constitutes an x-y projection of the 3D sperm movement.

Fig. 4 and 5 summarize the effects of emptying Ca^2+^ stores via the application of thapsigargin and cyclopiazonic acid. A total of 88 spermatozoa were analyzed for periods of up to 9.2 min, (4.6min and 9.2min bright field and fluorescence, as detailed in the Methods section: a. Bright field and fluorescence image acquisition). In order to contrast the behavior before and after applying the Ca^2^ modulators, we employed two controls (as described in Methods section: Spermatozoa samples and statistical analysis). The first one is provided by the same sperm 45 s before adding the inhibitors and its basal properties were compared to the rest of the time after inhibitor addition. Twenty-two spermatozoa were treated with thapsigargin and twenty-two with cyclopiazonic acid. The second control consisted of recordings without applying Ca^2+^ modulators to document the effect of long-time swimming itself, where we corroborated that these effects have no statistical significance (26 cells). In our experimental data set we found head/flagellar beat frequencies variations from 0.7 - 7 Hz / 6 Hz - 27 HZ respectively, with relative amplitudes increasing up to 4 fold from their control value.

**Fig. 4.**
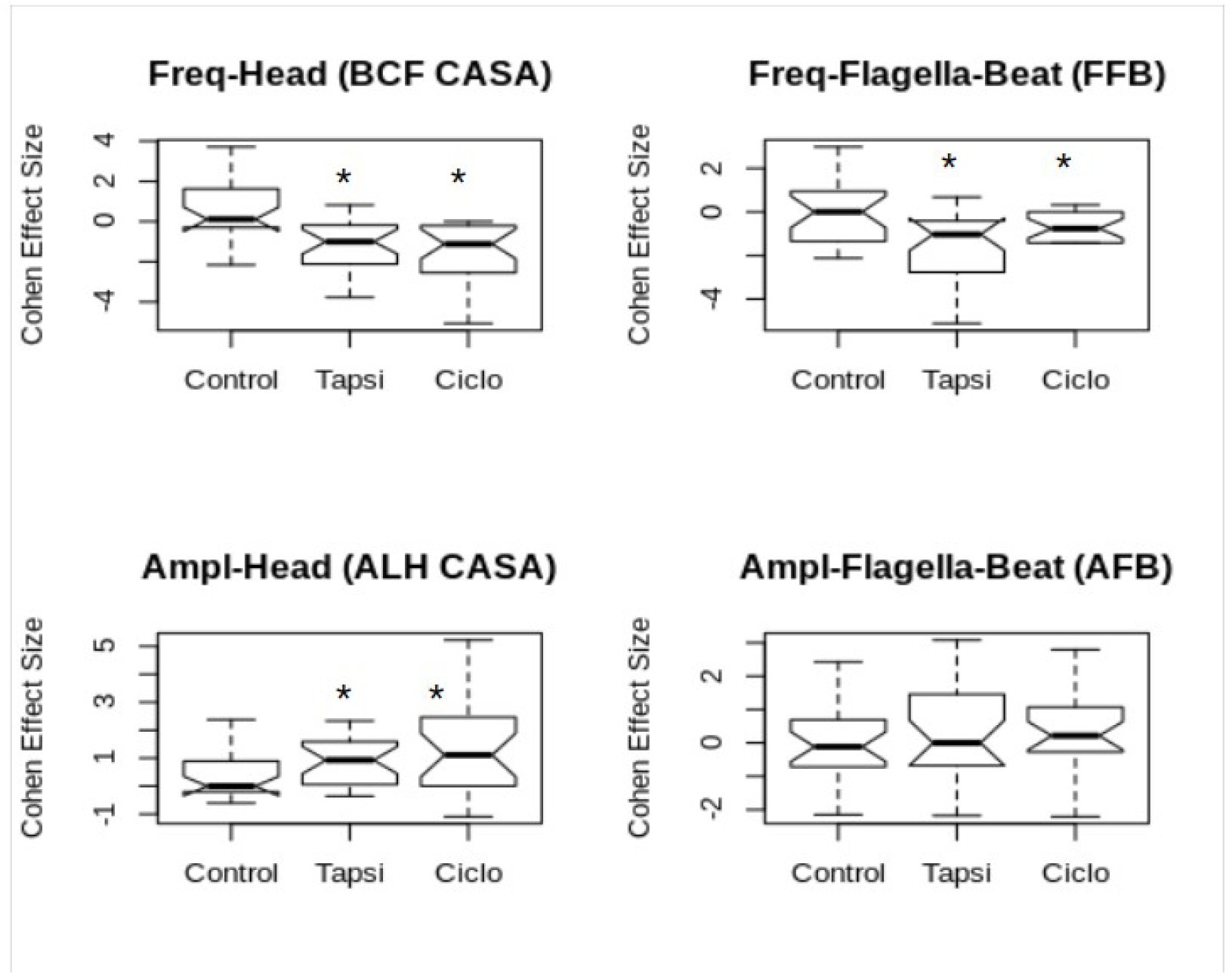
Emptying calcium stores with thapsigargin (22 cells) and cyclopiazonic acid (22 cells) produced statistically significant changes in three of the four kinematic parameters analyzed (Cohen Effect Size, Kruskall Wallys ANOVA, Eq. 1,2). Only the relative amplitude of the flagellar beat (AFB) did not show a statistically significant difference. a) Beat Cross Frequency (BCF CASA), b) Flagellar Frequency Beat (FFB), c) Relative Amplitude of Lateral Head displacement (ALH CASA), d) Relative Amplitude of Flagellar Beat (AFB).

**Fig. 5.**
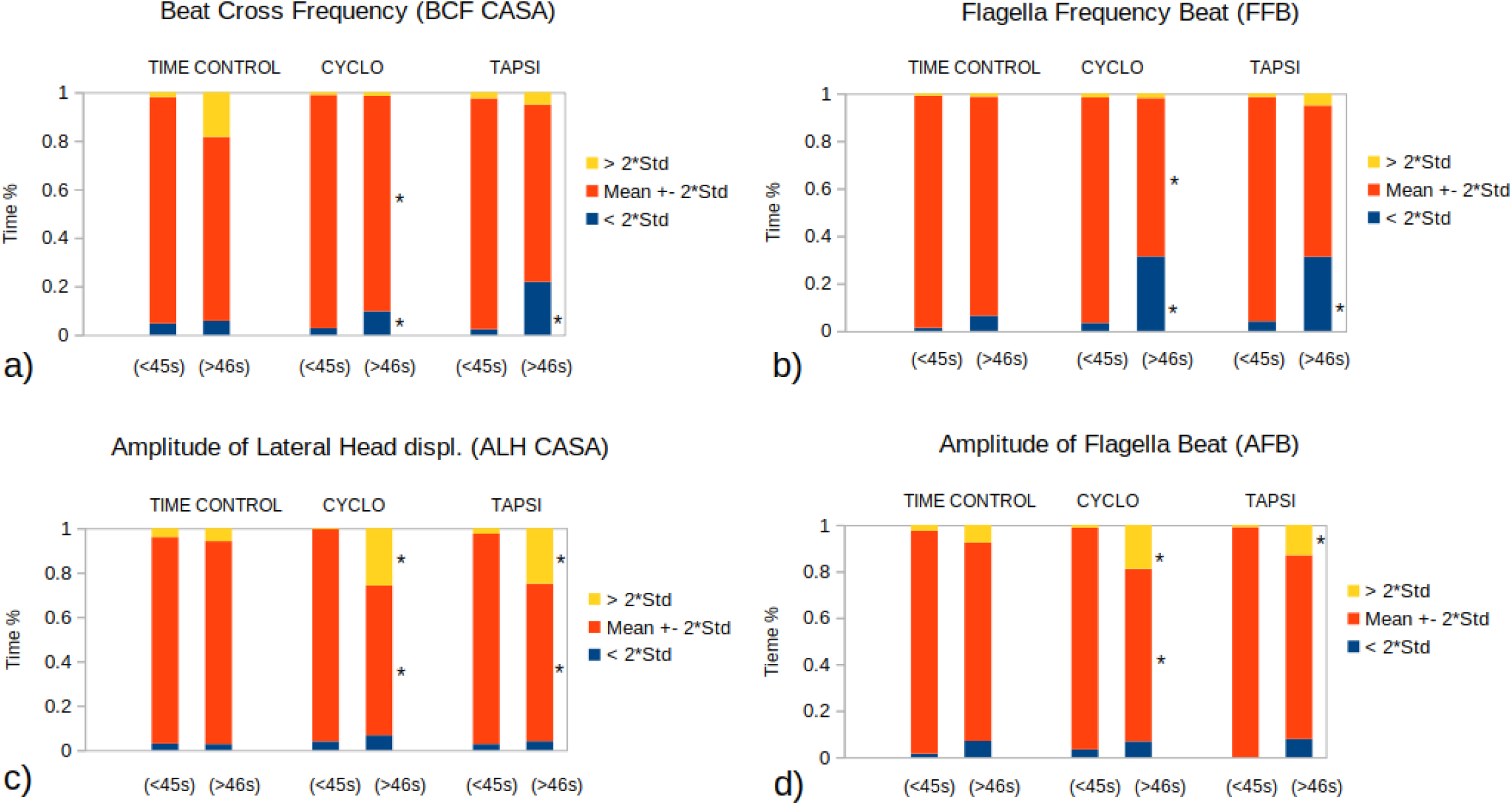
Total normalized swimming time of free-spermatozoa (for 4 kinematic evaluated parameters: BCF, FFB, ALH and AFB), in one of three determined categories: 1) In the control range (mean ± 2 std) (orange), 2) In the range 2 std above the control (yellow) and 3) In the range 2 std below the control (blue). The left bars for CYCLO and TAPSI pairs correspond to the first 45 s without applying calcium modulators (control) and the right bars for each pair to the rest of the time (up to 4.6 min) after their application. TIME CONTROL bars correspond to cells where no drugs were administrated. (*) represents statistical difference for α = 0.05. a) Beat Cross Frequency (BCF CASA), b) Flagellar Frequency Beat (FFB), c) Relative Amplitude of the Lateral Head displacement (ALH CASA) and d) Relative Amplitude of Flagellar Beat (AFB).

To contrast the differences before and after applying the Ca^2+^ modulators, we determined the Cohen’s *d* Effect Size for Anova and Kruskall Wallys for distributions that are not normal (see details in Methods section Eq. 1,2). Fig. 4 shows that emptying the Ca^2+^ stores has a significant effect in three of the four evaluated motility parameters, that is BCF, FFB and ALH (detailed in Table 1). Though increases were observed in the Amplitude of Flagellar Beat (AFB) after administration of thapsigargin and cyclopiazonic acid, though they were not statistically significant.

There is a large variance in spermatozoa swimming behavior, thus in Fig. 5 we show the relative total time when the cell swimming behavior is above or under two standard deviations (SD) of the control conditions. The yellow intervals correspond to the total percentage of time where the kinematic cell parameters are over two times SD and the blue bellow two times SD from control conditions (orange). The left columns from each pair correspond to the control condition before the application of cyclopiazonic acid or thapsigargin, while the right columns from each pair depict the effect after application of treatment.

It can be observed that the behavior during the control periods before treatment (Fig. 5 left columns for a, b, c, d) is quite symmetric: from the point of view that the areas above and bellow two standard deviations are similar (blue and yellow regions in Figs. 5). In contrast, when cyclopiazonic acid or thapsigargin are applied, the total time for both head yawing (BCF) and flagellar beating frequencies (FFB) decreased, as compared with control conditions (blue regions increase in Fig 5 for a, b right columns for treated cells). Inversely, the total time of the relative amplitudes of the head yawing (ALH) and flagellar beating (AFB) increased (yellow areas increase in Fig 5c, d right columns for treated cells). Significant statistical differences (using Wilcoxon test) for different ranges were marked with an asterisk in Fig 5. Notably, in this analysis the amplitude of the flagellar beating (AFB) increases in a statistically significant manner.

Fig 6 shows that after applying thapsigargin to empty the Ca^2+^ stores, sperm [Ca^2+^]i increases. This is in agreement with previous reports on [Ca^2+^]i (Meizel & Turner, 1993; Williams & Ford, 2003), and illustrates the capability of our methodology to record and accumulate statistics of [Ca^2+^]i during very long swimming periods.

**Fig. 6.**
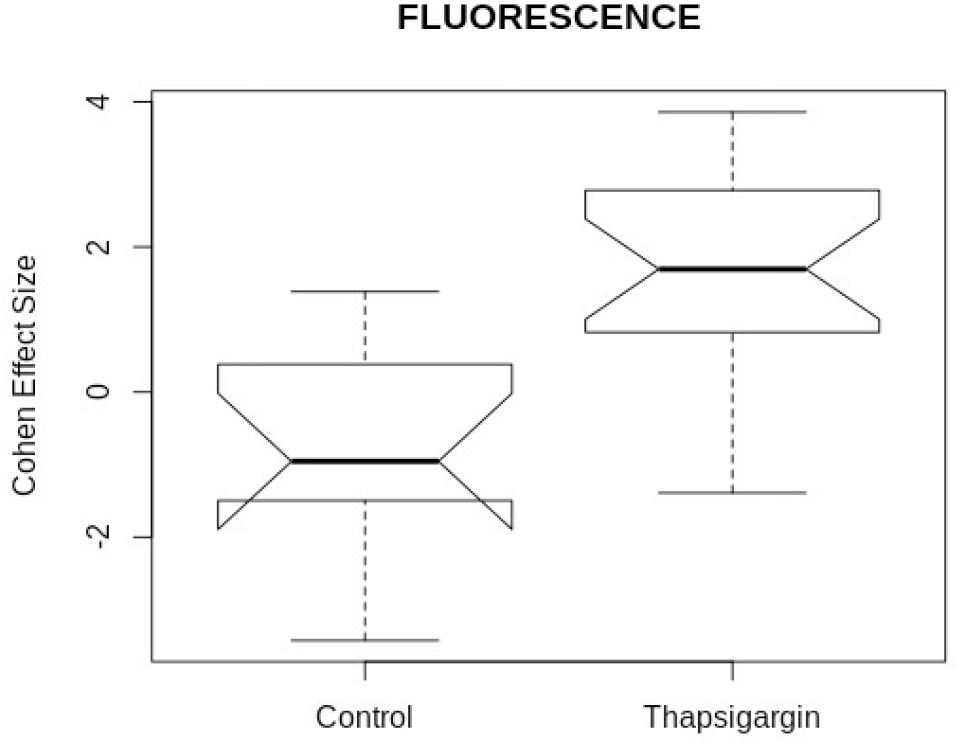
Thapsigargin addition increases [Ca^2+^]i.

### Comparison with segmentation-based sperm head and flagellar tracking systems

We have compared our results with those obtained with other available segmentation-based systems, such as SpermQ (Hansen et al., 2019) for flagellar and head kinematics, and OpenCASA (Alquézar-Baeta et al., 2019) for head kinematics only. Our microscope video recordings were not compatible with FAST (Gallagher et al., 2019) as it requires negative phase contrast microscopy. Due to uneven contrast, illumination, noise and existing debris in the background of our images, both SpermQ and OpenCASA had difficulties in tracking the sperm motion under these conditions, which in turn affected dramatically the processing time. SpermQ required more than 40 hours to process a single videomicroscopy recording containing 28,000 images, while our algorithm processed the same video data in less than 2 minutes. This demonstrates an important advantage of the developed segmentation-less procedure, which is able to robustly process non-ideal and noisy images with a fast processing time, especially relevant when working with videos with many thousands of image frames. SI Movie 4 illustrates how background noise and debris affected the segmentation output method in compared with the segmentation-less algorithm. Instead, the videomicroscopy data was filtered and pre-processed to allow extraction of kinematic parameters using SpermQ and OpenCASA, for comparison purposes, as displayed in Fig. 7.

**Fig. 7.**
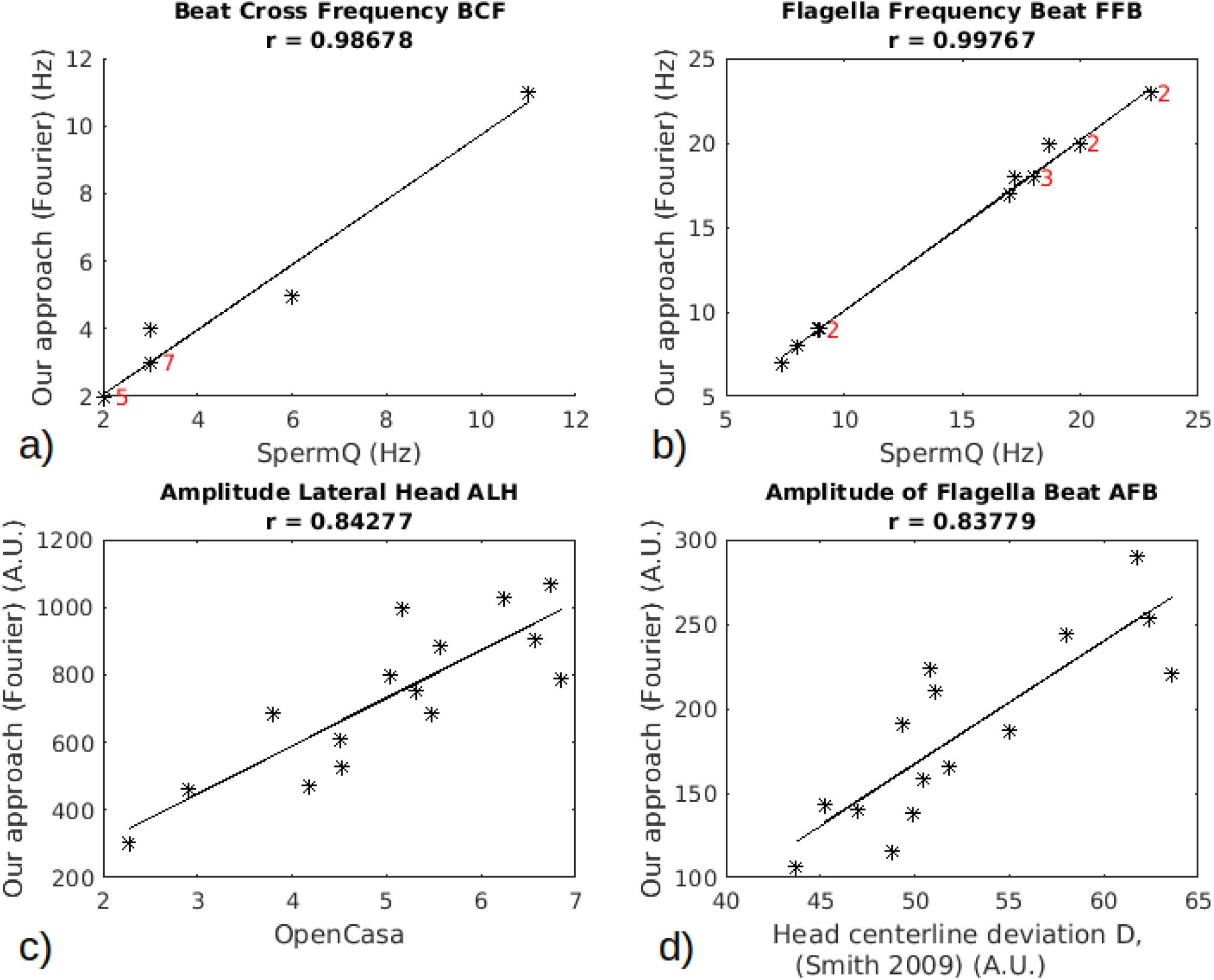
Comparison between the *segmentation-free* and segmentation-based methods showing strong correlations among all kinematic parameters for sixteen compared cells. The numbers inside a) and b) indicate the number of cells with the same value. a) Beat Cross Frequency BCF, compared with SpermQ (Hansen et al., 2019), Pearson’s *r* = 0.98; b) Flagellar Frequency Beat FFB, compared with SpermQ (Hansen et al., 2019), Pearson’s *r* = 0.99; c) Amplitude of Lateral Head displacement ALH, Compared with OpenCasa (Alquézar-Baeta et al., 2019), Pearson’s *r* = 0.84; d) Amplitude of Flagellar Beat AFB, compared with Head centerline deviation D defined by Smith, 2009, Pearson’s *r* = 0.84.

Fig. 7a & b display the high correlation for BCF and for FFB obtained between our method and SpermQ. The correlation values 0.99 and 1 respectively for the 15 cells evaluated. Regarding ALH, Mortimer et al., 2015 pointed out existing challenges in the different ways to evaluate this parameter, making it difficult to standardize between distinct algorithms in the literature. For example, inadequate smoothing of the average head trajectory may produce aberrant ALH values. Instead, we have compared our ALH results with those obtained with OpenCASA which is closely connected with our definition for this parameter. Fig 7c shows a strong correlation 0.84 with OpenCASA ALH. It is worth noting that OpenCASA found a correlation of 0.88 with respect to a commercial CASA system (Alquézar-Baeta et al., 2019).

Neither SpermQ, OpenCASA nor FAST (Hansen et al., 2019; Alquézar-Baeta et al., 2019; Gallagher et al., 2019) provide the amplitude of the flagellar beat AFB as an output, so we implemented the measurement of the ‘head centerline deviation’ D, as described in Smith et al., 2009 to measure this parameter. This allowed comparison between AFB measurement across all the different systems. Given that free-swimming sperm roll in 3D around their swimming direction (Gadêlha et al,. 2020), this introduces an oscillatory asymmetry in the ellipse-like head shape during cell swimming (SI movie 2). This biases distance measurements between the head main axis and the flagella in the perpendicular direction (Smith et al., 2009, Fig. 2a). Because of these issues, we used instead the mid-piece centerline which follows closely the head movement and eliminates noise from changes in head shape. We have found a strong correlation of 0.84 between both methodologies in Fig 7d for the head centerline deviation D (Smith et al., 2009, Fig. 2a).

## Discussion

The mechano-chemical micro-environment of the reproductive tract is complex. Spermatozoa must contend with dramatic physiological alterations in the ionic composition (Ng et al., 2018), temperature (Bedford, 2015), viscosity (Smith et al., 2009), fluid flow conditions (Gaffney et al., 2011; Ishimoto et al., 2017), as they undergo capacitation (Gervasi and Visconti, 2016) and may, in certain regions of the female reproductive tract, undergo rheotaxis (Miki & Clapham, 2013) and chemotaxis (Eisenbach and Giojolas, 2006) to be able to reach the site of fertilization (reviewed in Darszon et al., 2011, 2020). The spermatozoa’s ability to coordinate the flagellar beating over large distances and long swimming periods is thus critical for a successful fertilization. Against this background, the large majority of studies to date devoted to understanding sperm motility are critically limited to very short periods of observations. In this context, developing novel tools to record and process complex video microscopy imaging of sperm swimming for long time-periods is essential.

Computer Assisted Semen Analysis (CASA) technologies (Davis & Catz, 1996; Mortimer 2015; Gallagher 2018) derive primitive statistics of head trajectories by imaging samples over short periods of time e.g. 1 s, with a small field of view and for spermatozoa constrained to move in two-dimensions (CASA chambers have 10-20 microns depth). No direct flagellar kinematic parameter is assessed with CASA. High-precision spatio-temporal flagellar-tracking are equally constrained to short-time analysis (Smith et al., 2009; Hansen et al., 2019; Gallagher, et al., 2019; Ishimoto et al., 2017). One of the difficulties is that the flagellar waving occurs at a much faster rate (10-20 Hz) than sperm head lateral movement, requiring high-speed digital cameras (>100 fps) to resolve and track the flagellar beat. This requirement limits the total recording period to typically very few swimming strokes (Smith et al, 2009; Ishimoto et al., 2017). Even when flagellar tracking data is available, extraction of flagellar kinematic parameters, such as amplitude and frequency of the beat, are cumbersome in 2D (Hansen et al., 2019; Gallagher et al., 2019) and especially in 3D (Gadêlha et al., 2020).

An alternative to assess the long-time behavior of sperm beating is to pin the cell to the coverslip (Ooi et al., 2014; Saggiorato et al., 2017). This constrains the head position in the same field of view for imaging purposes. Nevertheless, the free-swimming behavior cannot be inferred directly from tethered sperm experiments (Gadêlha et al., 2020). The flagellar beat of a pinned spermatozoon is modulated differently from a free-swimming one (*ibid*). Human spermatozoa roll around the swimming direction via a complex interplay between one-sided asymmetric strokes and out-of-plane pulsations, responsible for the sperm rolling in 3D (Gadêlha et al., 2020), whilst pinned sperm switch their beating to a planar and symmetric side-to-side movement (Saggiorato et al. 2017; Ooi et al., 2014). It is thus unclear how to best characterize the motility of free-swimming spermatozoa over long periods, especially for durations commensurate with the time required for sperm accession in the female reproductive tract. Indeed, swimming patterns and long-time behavior of freely swimming human spermatozoa remain elusive in the literature (Achikanu et al., 2019). Are, for example, swimming patterns observed over short-periods representative of sub-populations with distinct motility traits, or instead, individual sperm are capable of generating multiple swimming gaits when observed over long-periods? Furthermore, what is the relation between the motility information measured from short-time experiments with their long-period counterpart? All these questions are now receiving more attention as the multifaceted ability of spermatozoa to adapt the flagellar behavior over the course of long periods of time is unveiled (Achikanu et al., 2019).

Bioimaging segmentation is at the heart of any automated computer-assisted image analysis system. The most recent systems implementing a range of image segmentation strategies include SpermQ (Hansen et al., 2019), FAST (Gallagher et al., 2019) and OpenCASA (Alquézar-Baeta et al., 2019). However, segmentation-based methods are based on thresholding the pixel intensity of the image, thus sensitive to the imaging quality, background noise, defocusing and illumination effects. This sensitivity limits the general applicability of thresholding methods, often requiring high-level of algorithmic customization to fit the specific needs of each bio-imaging data.

Here, we solved a critical bottleneck in computer-assisted flagellar motility analysis. We developed (1) a *segmentation-free* analysis system which in turns allows for the (2) inspection of flagellar beating-pattern of individual cells over long periods of time (up to 9.2 min). The sperm beating can thus be analyzed over tens of thousands of flagellar beat cycles, as required during long swimming distances within the female reproductive tract. The *segmentation-free* analysis is based on the local orientation and anisotropic features of an image (Püspöki et al., 2016), and can be easily extended to general videomicroscopy imaging data with any duration of other flagellated microorganisms, or indeed slender-body organisms, such as *Chlamydomonas*, trypanosome and *c. elegans*. The method provides a temporal series representing the orientation of the swimming sperm at each time-frame. As such, the orientation reduces the complexity and dimensionality of the data to a one-dimensional signal (a function of time only) and embodies both head and flagellum kinematic parameters. High-dimensional data sets are intrinsic to flagellar shape analysis (Werner et al., 2014). Both head and flagellar kinematics parameters can be directly disentangled from the one-dimensional orientation signal using simple Fourier decomposition. This bypass convoluted spatial-temporal flagellar tracking and analysis (Gadêlha et al., 2020; Werner et al., 2014). Thus in addition to the head-trajectories and swimming directionality, the frequency and amplitude of both head lateral oscillation and flagellar waving movement are also obtained with this strategy.

Using the computer-assisted *segmentation-free* strategy, we investigated the role of emptying calcium stores during long swimming periods in human spermatozoa. In many cellular systems, the release of Ca^2+^ from internal stores is known to contribute significantly to the overall concentration of [Ca^2+^]i and its changes, thus driving specific signaling responses (Clapham et al., 2007; Putney et al., 2013). Changes in the sperm [Ca^2+^]i are implicated in the regulation of flagellar beating (Darszon et al., 2011; Strünker et al., 2015; Lishko & Mannowetz, 2018), and they are influenced by Ca^2+^ release and re-uptake from internal stores such as the acrosome and the residual nuclear vesicles (Chang & Suarez, 2011; Correia et al., 2015). Ion channels, for instance the IP_3_ Receptor (Prole & Taylor, 2019) and the Ryanodine Receptor (Ogawa et al., 2020), able to release Ca^2+^ from internal stores, have been located to both of these sperm internal stores (reviewed in Correia et al., 2015). In many somatic cell types, store Ca^2+^ mobilization is triggered by intracellular messengers like cADPR, NAADP and IP_3_ (Galione & Chuang, 2020). This has been documented to occur in mammalian sperm (Correia et al., 2015). Ca^2+^ release can also be induced by inhibiting the Ca^2+^-ATPases of these stores, as we have done in the present work, or by Ca^2+^ waves triggered by external ligands, as progesterone in human sperm (Blackmore et al., 1999; Garcia & Meizel, 1999; Harper et al., 2004). As [Ca^2+^]i increases in the vicinity of these stores, it triggers what is known as Ca^2+^ induced Ca^2+^ increases by activating RyRs and IP_3_Rs (Parkash & Asotra, 2012). It had been shown previously that releasing Ca^2+^ from internal stores with thimerosal, an activator of IP_3_R and RyR channels (Elfering et al., 1999), induced hyperactivation in mouse and human sperm. Notably, thimerosal could induce hyperactivation in the absence of external Ca^2+^ clearly indicating the effect was due to Ca^2+^ release from internal stores (Marquez et al., 2007; Alasmari et al., 2013). Thapsigargin has been used in several mammalian sperm to induces Ca^2+^ release from internal stores, in many of them it is able to induce hyperactivation and in some the acrosome reaction (reviewed in Correia et al., 2015).

Here we found that elevating [Ca^2+^]i by emptying the human sperm internal Ca^2+^ stores with two different store Ca^2+^-ATPase inhibitors, thapsigargin and cyclopiazonic acid, significantly decreases head and flagella beating frequencies and increases the head beat amplitude in a statistically significant manner. As would be expected for hyperactivation inducers, the amplitude of the flagellar beat also increases but without reaching significance. However, this latter parameter does significantly increase when the relative total time that this parameter spends above two standard deviations of the control condition is determined. These findings are in agreement with most previous reports of hyperactivation stimulatory conditions mentioned earlier and reported in the literature (Ho & Suarez, 2001, 2003; Chang & Suarez, 2011; Correia et al., 2015). There are published exceptions such as Rossato et al., 2001, where 10 μM thapsigargin and up to 100 μM cyclopiazonic do not affect motility (see also Vijayaraghavan et al., 1994; Williams & Ford 2003). It is worth pointing out that few studies have been reported with cyclopiazonic acid. Our findings highlight the usefulness of our new segmentation-free strategy to study and correlate sperm swimming properties and [Ca^2+^]i, a matter that is fundamental and still requires further research and analysis.

## Conclusions

CASA measurements are typically performed in short intervals of time, i.e. 1 s. from where kinematic parameters are estimated. This way of sampling assumes that sperm behaviour is stable for long periods of time. Recent evidence indicates this is not true, kinematic parameters are continuously changing and dynamically adapting. In this work we present a novel methodology based on the local orientation and isotropy of individual free-swimming spermatozoa without the need of bio-image segmentation (thresholding). This in turns allows for extraction of long-term kinematic and physiological parameters (up to 9.2 min). We have demonstrated that the spermatozoa head and tail beat may have important variations in frequency and amplitude over time. Furthermore, our strategy is compatible with fluorescent recordings to follow the associated [Ca^2+^]i changes that occur upon sperm switching between different swimming modes.

Particularly, with this methodology we were able to evaluate the frequency and relative amplitude of the lateral head displacement (BCF and ALH in CASA and OpenCASA standards) and of the flagellar beating (FFB and AFB) without the need of segmentation procedures. We could observe and measure significant effects when emptying calcium stores, particularly in the head and flagellar beating frequencies and relative amplitudes for human spermatozoa. This was possible as the strategy permitted taking short samples (avoiding cell damaging) of a [Ca^2+^]i sensitive dye emitted fluorescence for the whole long-term evaluation before and after applying these drugs to the same spermatozoon. The *segmentation-free* procedure presented significant advantages as compared with segmentation-based procedures. The main advantages of the proposed methodology are, besides avoiding the substantial effort needed to segment the flagella, the insensitivity to noisy background and artifacts in the images, as well as an important tolerance to defocusing and heterogeneous background illumination, and fast processing times of large data (less than 2min). The estimated parameters obtained with this methodology were compared with those estimated with other systems observing high correlations. Notably, this new strategy may be applied to other swimming microorganisms, such as spermatozoa of any species, *Chlamydomonas (*Polin et al., 2009), trypanosomes (Gadêlha 2007), and even *c. elegans (*Ding et al., 2019), for example, by exploiting their inherit slender-body shape which further defines an overall orientation during motion. Thus, this method may appeal to distant research communities away from sperm physiology and reproductive biology.

## Supporting Information

### a. Spermatozoa orientation detection and tracking

Orientation detection from an image is important for a wide range of applications. A popular method to extract orientation relies on gradient information at each position **x**=(x0,y0) in the image, where the first-order directional derivatives are computed. The directional derivatives become low when the direction is similar to the orientation at point **x**. Orientation calculation using this approach is popular because it is easy to implement and provides good approximations. OrientationJ (Püspöki Z., 2016, R. Rezakhaniha, 2012, E. Fonck, 2019) is a FIJI plugin (Schindelin, 2012) developed to more efficiently compute the average orientation of a region of interest (ROI) in an image. This software employs a more robust approach (less sensitive to noise) based on the use of a tensor structure given by

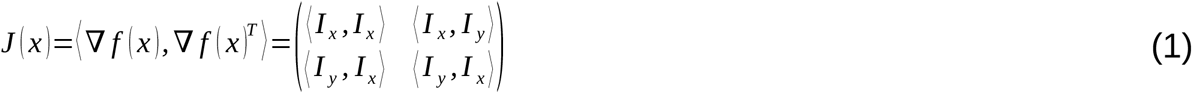

where the image I is convolved with a Gaussian filter to remove noise, the eigenvalues of matrix J(x) provide local shape information (E. Fonck, 2009; Harris C., 1988) and cubic B-spline interpolation is used to compute the continuous spatial derivatives. The two eigenvalues of J(**x**) give local shape information in the neighborhood of point **x**. If the two eigenvalues are similar and close to zero then, the region is homogeneous (noisy structure), if the two eigenvalues are similar and larger than zero then the region is rotationally symmetric (blob structure), if one is positive and large and the other is close to zero, then the eigenvalue is aligned with the gradient direction (line structure). The eigenvector associated to the largest eigenvalue gives the dominant direction. It can be shown that the orientation of the dominant direction at a given position **x** can be computed by:

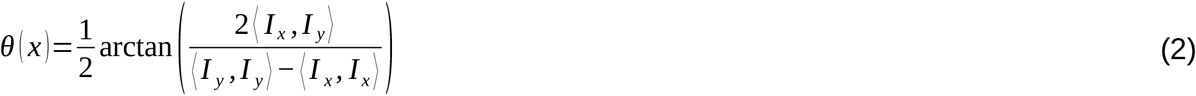

To compute the orientation over a ROI, the derivatives are summed over the ROI

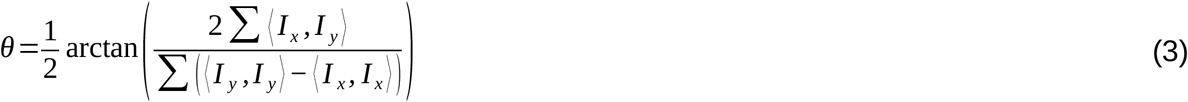

This approach to compute the dominant direction on the ROI performs very well and therefore it is feasible to apply it to obtain the flagellum orientation in an image. Nevertheless, for free moving cells, it is difficult to have clear images with only one single cell in the field of view, thus the computation of the flagellum orientation (the derivatives are summed over the ROI, usually the full image) gets incorrectly affected by other cells or artifacts. For that reason, we modified and extended the functionality of OrientationJ to a more robust dominant spermatozoa flagellum orientation detector avoiding adding orientations from spurious objects. This was achieved by tracking the spermatozoa and measuring locally its main dominant orientation.

### b. Image stack pre-processing

The image sequences (bright field microscopy) need to be firstly pre-processed. This was done with standard Fiji’s commands as follows: a) Project the original image stack along the axis perpendicular to image plane (Z Project with average intensity). The resultant projection will contain not moving artifacts and the background (even or uneven) illumination. b) Invert both original stack and its corresponding averaged Z Projection. c) Subtract from original stack the averaged Z projection to eliminate not moving artifacts and uneven background illumination. d) Enhance the contrast by adjusting the brightness/contrast (use the automatic mode), and apply the improved look-up tables. At this point the spermatozoon flagellum is practically binary, white over black background. e) Apply a median filter with a radius kernel of 2 or 4 pixel to eliminate impulsive noise and artifacts. At this point, the stack is ready to be processed by the modified Orientation J plugin.

### c. Modified OrientationJ plugin

The user has to identify the spermatozoon of interest by clicking over the head to determine the initial position **p0** for tracking. Then, the dominant direction (Eq. (2)) is calculated using three different ROIs: (i) a squared ROI with center at **p0** and size WS is employed to calculate the spermatozoon orientation **O**_square_. The square region must be big enough to contain the spermatozoon head and flagellum information. Note that the larger the square, the more plausible the orientation can be affected by external objects; (ii) a circular ROI with center at **p0** and radius **r0** is employed to compute the spermatozoon head orientation Ocircle. The radius **r0** must be big enough to cover the spermatozoon head without including the spermatozoon flagellum and (iii) a rectangle ROI with height **h**, width **w** and rectangle width parallel to direction **O**_square_ is constructed to compute the spermatozoon flagellum orientation, the points belonging to circular ROI are excluded from the analysis. Hence, this region should include orientations only from the spermatozoon flagellum eliminating the influence of orientations coming from the spermatozoon head. Note that the larger the width, the more flagellum information can be included to compute the flagellum orientation. Fig. 1 depicts the three different ROIs employed to compute the dominant direction. Rectangular ROIs should be used for analysis, however this is sometimes unfeasible because the flagellum gets out of focus and there is no dominant direction. In our analysis, we use square ROIs because they are more stable. In addition to measuring the dominant direction, we also keep track of three measures: mean intensity, coherence (Püspöki Z., 2016) and magnitude of dominant direction for the square ROI. These measures are useful to identify if the spermatozoon head is still being tracked (we may have cases that the track does not follows the spermatozoon head because it is out of the field of view in the XY plane or it is moving too much in the z-direction). Once the spermatozoon orientation has been computed for time **t**, we proceed to the following time-point by automatically updating the spermatozoon position **p**t+1. To achieve this, a weighted sum over the positions inside the circle ROI and image intensity is calculated by:

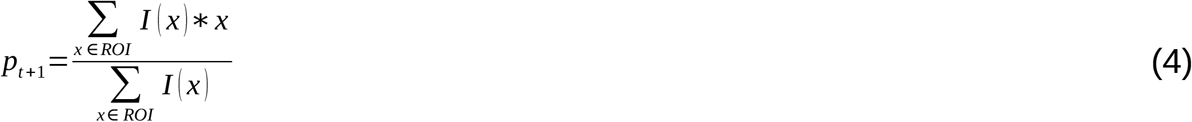

where x is a point (two coordinates) in the image and I(x) is the image intensity at position x. This approach places **p** t+1 near the spermatozoon head center at time **t+1**. However, this process can be affected if the spermatozoon moved a larger distance from the circle ROI at time **t+1** or if it disappears from the image or there is another spermatozoon crossing the circle ROI. In such cases, the spermatozoon track can be recovered at the current time **t+1** or a posterior times if it has disappeared. We proposed two conditions to verify that a square ROI contains the spermatozoon head: (i) the first condition takes into account the coherency, it must be larger than 0.1 and; (ii) the second condition takes into account the square mean intensity over the ROI, it must not change by a large amount from previously identified spermatozoa. This implies that the mean intensity over the square ROI for time **t+1** must be larger than 0.4* *μ*_*I*_ where *μ*_*I*_ is the mean the mean of the measure mean intensity over previous times that were identify as correctly having a spermatozoon head. The value 0.4 was found experimentally. If these two conditions are satisfied, a spermatozoon exists in the current square ROI, otherwise it is inexistent. In the case that there is no spermatozoon, then the full image is covered with not overlapping square ROIs. The square ROI with the lowest cost is identified as the best ROI to compute the orientation at the current time. The cost for each square ROI is given by

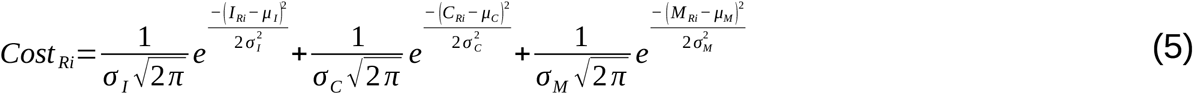

where *R*_*i*_ is the current region to analyze, 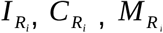 are the mean intensity, coherency and magnitude over region RI (measures of region), respectively. *μ*_*I*_,*μ*_*C*_ and *μ*_*M*_are the mean intensity, mean coherency and mean magnitude of the measures from regions of previous times that have been identified as correctly including a spermatozoon head. *σ_I_*, *σ_C_*and *σ_M_*, are the standard deviations from the Gaussian distribution and were set experimentally to 10, 0.5, 10. The cost is designed to give highest priority to regions having similar measures than the previous detected regions. Once that the best square ROI is selected for time **t+1**, the algorithm computes the three orientation measures presented previously and continue with time **t+2**, and the process is repeated until no images are left.

Our approach to compute the orientation has three main advantages over the original OrientationJ version (Schindelin, 2012): (i) it is less sensitive to external objects since it is computed over a neighborhood from the spermatozoon, (ii) it is fast since the tensor structure J(x) has to be computed only on a small squared region (except for the few cases that the algorithm missed the spermatozoon and has to re-compute the best ROI) instead of computing the tensor over the full image and (iii) our approach is customized to obtain three different orientations which can be better suited for the user needs than a single orientation.

